# *APOE* genotype and sex drive microbiome divergence after microbiome standardization in *APOE*-humanized mice

**DOI:** 10.1101/2025.04.25.650741

**Authors:** Michelle Aries Marchington, Hope Gasvoda, Makayla Michelotti, Fernando Rodriguez-Caro, Ashley Gooman, Anna Perez, Tiffany Hensley-McBain

## Abstract

The *APOE4* allele is the greatest known genetic factor for sporadic or late onset Alzheimer’s Disease (LOAD). Gut microbiome (GMB) dysbiosis can lead to poorer outcomes in disease. The intersection of sex, *APOE* genotype, inflammation, and gut microbiota is incompletely understood. Previous studies in humans and humanized *APOE* mice have demonstrated *APOE*-genotype specific differences in the GMB. However, most of these studies were unable to resolve bacteria to the species level. It remains unclear how GMB changes with age and sex in the context of *APOE* genotype. In this study, humanized male mice with either *APOE* 2, 3, or 4 genotype were bred with the same two C57BL/6J sisters to standardize microbiomes across lines and monitor divergence based on *APOE* allele. Stool samples were collected at breeder set up and from the heterozygous (F1) and homozygous (F2) generations at wean and 6 months old. Stool was assessed via shallow shotgun sequencing to enable species and strain level taxonomic resolution. The heterozygous pups’ microbiome resembled each other at wean across all genotypes. However, the heterozygous pups, and their homozygous offspring continued to diverge, particularly the *APOE2* females. In homozygous mice, the GMB demonstrated significance divergence at 6 months of age based on sex and *APOE* genotype. In comparison to their *APOE3* and *APOE4* counterparts, *APOE2* females and males demonstrated an increased quantity of bacteria associated with anti-inflammatory profiles, including in the *Lachnospiraceae* family *(Lachnospiraceae bacterium UBA3401*) and decreased quantities in the *Turicibacteraceae* family (higher levels are associated with LOAD).

**Importance:** The *APOE4* allele is implicated as a significant risk factor for many diseases including cardiovascular disease (responsible for more deaths than any other disease) and sporadic or late onset Alzheimer’s Disease (accounts for an estimated 60% to 80% of all dementia cases). It is known that the gut microbiome (GMB) is affected by different genotypes and disease states. Mouse model studies have environmental and genetic controls allowing a specific gene to be studied. This study aims at discovering key GMB species differences allowing for future therapeutic targets. The GMB of the experimental mice was standardized and genotype and sex-specific divergence was observed with species and even strain level taxonomic resolution. Reported here are the first data demonstrating GMB divergence over time driven by *APOE* genotype from an inherited source and the first data to identify APOE genotype-specific bacteria species that may serve as therapeutic targets in *APOE*-driven disease.

## Introduction

Apolipoprotein E (ApoE) is a lipid cargo binding protein found in high quantities in the brain that is produced by and interacts with immune cells, has been shown to impact the gut microbiome, and is implicated in several diseases^1,2,3^. In humans, there are 3 major isoforms of ApoE: ApoE2, ApoE3, and ApoE4 corresponding to 3 main alleles in the population (*APOE2, APOE3*, and *APOE4*). *APOE3* is the most common, with *APOE2* and *APOE4* each containing a single point mutation resulting in one amino acid change from ApoE3 isoform^4^. Despite the single amino change between isoforms, different *APOE* genotypes have been implicated in a multitude of altered biological functions, including altered immune and lipid binding and altered signaling, and associates with increased risk of multiple diseases such as Alzheimer’s disease (AD) and cardiovascular disease (CVD)^3,5,6,7^.

AD is a debilitating disease that affects approximately 6.8 million persons in the United States, two-thirds of which are women, and accounts for an estimated 60% to 80% of dementia cases^8^. AD is characterized by the accumulation amyloid beta plaques and neurofibrillary tau tangles, which can lead to a variety of cognitive symptoms, including muscle control loss, behavioral changes, and memory loss. Age is the greatest risk factor for AD, but genetic and environmental factors also play a role^3,9^. *APOE4* is the greatest known genetic risk factor for sporadic, or late-onset Alzheimer’s Disease (LOAD), and individuals homozygous for *APOE4* have a 12-fold increased risk of AD when compared to individuals homozygous for *APOE3*^3,9^. Isoform-specific differences in binding affinity to immune receptors and lipids have been implicated in differential disease risk^9,10^. *APOE4* carriers with AD show earlier Aβ accumulation, earlier clinical disease onset, faster disease progression, heavier plaque burden, and increased brain atrophy, while *APOE2* carriers with AD have later Aβ deposition, later clinical onset, and increased longevity^1^.

CVD is an ever-increasing health concern that is responsible for more deaths worldwide than any other disease (13 %)^11^. The link between *APOE* genotype and CVD has been attributed in part to altered binding affinities and lipid transport resulting in altered cholesterol, HDL, and LDL levels^10^. Moreover, it is known that the gut microbiome plays a role in CVD, including the association between circulating Trimethylamine-N-oxide, a gut microbiota-derived plasma metabolite, and an increased risk of CVD and cardiac related events^12^. The circulation of this metabolite, which is also elevated in AD, results in an inflammatory response and increases vascular inflammation^12,13^.

The intersection of sex, *APOE* genotyp*e*, inflammation, and gut microbiota (GMB) is incompletely understood. It is known that genotype can drive GMB changes, and that the GMB is a key essential to health. Several previous studies have suggested that *APOE* genotype is associated with different GMB profiles, inflammation, and different disease states^14^. However, mechanistic studies on *APOE* genotype as a driver of microbiome change and studies that can identify taxa to the species level are lacking. Hammond et al. identified genotype dependent differences in commensal bacterial species in humans 55-85 years of age with different *APOE* genotypes^15^, and several other human studies have identified family or genus level changes^2,16,17^. Previous mouse studies assessing *APOE* and the GMB used 16s rRNA to identify the microbiota down to the family level and did not use mice standardized for microbiomes across lineages to assess the true impact of genotype^2,16,18,19^. Species level resolution is needed to identify which bacteria are driving inflammation and determine immune microbial therapeutic targets, as different bacteria within the same family can possess different inflammatory profiles.

Here, we present a study investigating the role of *APOE* genotype in driving GMB divergence in a humanized targeted replacement mice with either *APOE* 2, 3, or 4, allowing for a controlled environment to assess genotype-specific impacts without external factors that impact the human GMB (diet, exercise, environment, etc). It is known that animals with independent lineages even within the same facility demonstrate different compositions of gut microbiota^20,21^, and one method of microbiome standardization includes back-crossing mice of different lineages or genotypes back to the same mother^20^. In this study, we bred *APOE* 2, 3 and 4, mice separately back to the same C57BL/6J sisters to investigate how a standardized and inherited GMB changes with age and sex in the context of *APOE* genotype and to identify *APOE* genotype-specific bacterial species that may be targets for microbiome modulation in *APOE*-driven disease.

## Methods

### Animals

Four mouse strains were used in this study: 1) humanized *APOE2* (B6.Cg-Apoe<em3(APOE*)Adiuj>/J (Strain: 029017)), humanized *APOE3* (B6.Cg-Apoe<em2(APOE*)Adiuj>/J (Strain: 029018)), humanized *APOE4* (B6(SJL)-Apoe<tm1.1(APOE*4)Adiuj>/J (Strain: 027894)), and in-house maintained C57BL/6J (B6) (Strain: 000664). The in-house maintained lines are periodically refreshed from The Jackson Laboratory (JAX) to decrease genetic drift. *APOE* 2, 3, and 4 mice were purchased from JAX. Mice were housed in individually ventilated and air-filtered cages in a super-barrier mouse room. All cages, bedding, food, water, and enrichment materials were autoclaved or UV-treated prior to contact with the mice. All cages were opened only in a biosafety cabinet, and personnel wore autoclaved lab coats and sterile gloves. Mice had free access to food, water, and enrichment materials. All Institutional Animal Care and Use Committee (IACUC) protocols were followed. All research followed *The Guide for the Care and Use of Laboratory Animals, and the Public Health Service Policy on Humane Care and Use of Laboratory Animals*.

### Microbiome Standardization

Two B6 female litter mates were bred first to an *APOE4* male, then to an *APOE2* male, then to an *APOE3* male, resulting in offspring heterozygous for the human *APOE* 2, 3, or 4 gene (N1 generation) (Fig. 1). Heterozygous N1 mice of the same genotype were bred together, and homozygous pups were selected to generate standardized microbiome lines for each genotype (N1F1 and following generations) (Fig. 1). To account for cage-to-cage effects samples collected for microbiome assessment for each genotype were collected across multiple cages of mice.

**Figure 1.**
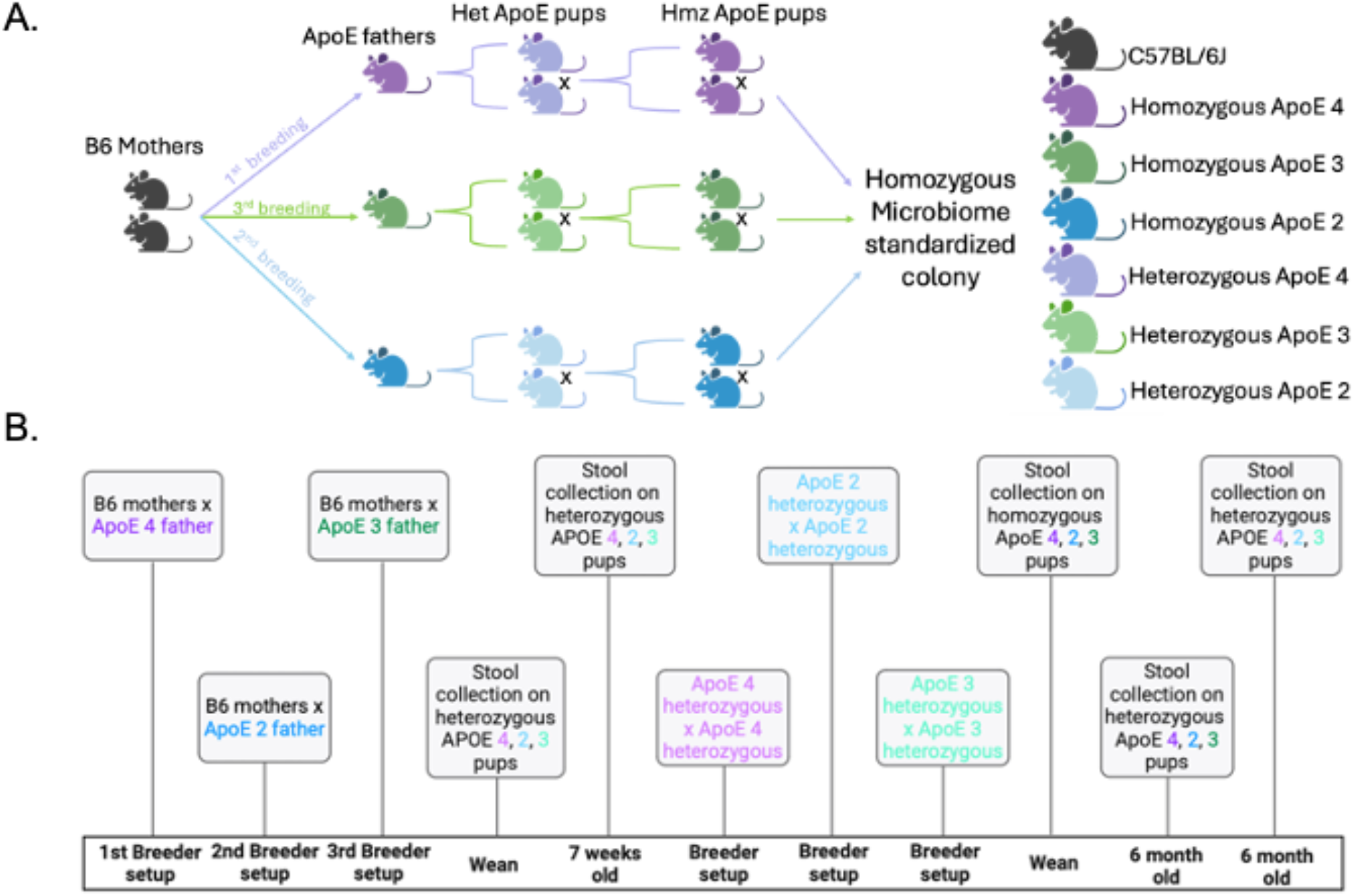
Breeding schematic and timeline. A. Breeding schematic for the generation of a microbiome standardized homozygous humanized *APOE* 2,3, and 4 mouse colony. B. Timeline on stool collection of breeder and aged pups of N1 and N1F2 generation.

### Stool Collection

Fecal pellets were collected from the B6 mothers and *APOE* 2, 3, and 4 fathers at breeder setup (8-weeks-old). Fecal pellets were collected from the N1 heterozygous offspring at wean, 7 weeks old, and 6 months old and from the homozygous offspring (N1F2 generation) at wean and 6 months old. All personnel wore autoclaved lab coats and sterile gloves for stool collection. Mouse cages were opened only in a biosafety cabinet. Mice were placed individually into empty autoclaved cages for the duration of stool collection to ensure proper identification of each sample. Stool pellets were placed into sterile 2 mL screwcap tubes and stored at −80°C until shipment to Transnetyx and transferred to room-temperature stable storage buffer (Transnetyx Microbiome Sample Collection Kit) prior to shipment.

### Shallow Shotgun Sequencing

Shallow shotgun sequencing was used to increase the sequencing depth and enable species and strain level taxonomic resolution. Stool was shipped in a DNA stabilization buffer to Transnetyx for extraction, library preparation, and sequencing. Genomic DNA was converted to sequencing libraries with unique dual indexed adapters to ensure that reads and/or organisms are not misaligned (Transnetyx Inc., Cordova, TN, USA). DNA extraction is performed using the Qiagen DNeasy 96 PowerSoil Pro QIAcube HT extraction kit and protocol. Genomic DNA is converted to sequencing libraries with unique dual indexed adapters to ensure that reads and/or organisms are not misaligned. Library preparation is performed using the Watchmaker DNA library preparation with fragmentation protocol. Sequencing is performed using the Illumina NovaSeq instrument and protocol at a depth of 2 million 2 × 150 bp read pairs. Raw data (in the form of FASTQ files) was analyzed using the One Codex analysis software using the One Codex database consisting of more than 127,000 whole microbial reference genomes^22^. Comparing a microbial sample against the One Codex Database consists of three sequential steps: 1) Every individual sequence is compared against the database by exact alignment using k-mers where k=31^23,24^; 2) Sequence artifacts are filtered out based on the relative frequency of unique k-mers; 3) The relative abundance of each microbial species is estimated based on the depth and coverage of sequencing across every available reference genome.

### Filtering

Data was processed using R version 4.3.2. To account for differences in library size across samples, data was subsampled and standardized using the microbiomeStat (v1.2.0) R package. First, read counts assigned to the host species (*M. musculus*) or to unannotated microbial species were removed. Next, samples were rarefied using a maximum threshold of three million read counts and then standardized using the Total Sums Scale method. Microbial species were filtered using an abundance threshold of at least 0.005% within each timepoint and zygosity group and filtered to include only species present in at least 20% of the total number of male or female samples. A three million reads threshold was selected for rarefaction to equalize statistical power across all timepoints due to differences in library sizes across samples. Batch effects after rarefaction was tested by adding batch as a variable in the PERMANOVA models before and after standardization and confirmed no batch effects after rarefaction and consistency of results across the subsampling threshold.

### Statistical analysis

Clustering of samples was evaluated through principal component analysis (PCA) using the prcomp function of the “base” R package. Community composition differences were tested through PERMANOVA tests using the adonis function from the “vegan” R package (v. 2.6.8) and selecting the “Bray-Curtis” method to calculate pairwise distances. Microbial signatures were estimated using the coda_glmnet function from the “coda4microbiome” R package (v.0.2.4). The Galaxy Metabiome Portal was used to conduct a Linear discriminate analysis Effect Size (LEfSe) and Cladogram representing data from the LEfSe on the 6-month-old homozygous and heterozygous mice to compare the GMB among humanized *APOE* 2,3 and 4 mice^25^. One Codex’s platform was used to generate bacteria species abundance and Shannon diversity plots. Dot plots with a 2-way ANOVA followed by Tukey’s multiple comparisons test with multiplicity adjusted p-values were generated using Prism 10 version 10.1.1.

## Results

The GMB of heterozygous *APOE2, APOE3*, and *APOE4* mice resembled that of each other and that of the B6 mothers at wean (Fig. 2A). The female pups clustered together based on GMB composition at wean (Fig. 2B) with no statistically significant differences in GMB composition (p>0.1 by PERMANOVA). However, male pups’ GMB composition demonstrated some divergence at wean (Fig. 2C) with a statistically significant difference driven mainly by the *APOE4* pups (p<0.01 by PERMANOVA). Heterozygous *APOE* 2,3, and 4 pups show nonconformity and no reversion to the microbiomes of their fathers (purchased homozygous *APOE* 2, 3 and 4 males) from wean to 6-months-old (Sup. Figs. 1B and 2A-B). Genotype based divergence of the heterozygous pups was observed by 6-months-old (Sup. Figs. 2 and 3). PERMANOVA comparing sex-specific GMB changes in *APOE* zygosity 6-month-old mice, showed no statistically significant differences between homozygous or heterozygous male mice having an *APOE4* allele (p<0.3), or having an *APOE3* allele (p<0.1), while *APOE2* mice had a statistically significant difference based on zygosity (p<0.05). When comparing zygosity in 6-month-old female mice, *APOE*4 mice demonstrated significant differences between homozygous and heterozygous mice (p<0.05), while *APOE*3 mice trended toward significance based on zygosity (p=0.051), and *APOE*2 mice were not significantly different by PERMANOVA. The heterozygous mice were bred to generate the homozygous experimental line and the GMB was tracked to 6 months of age (Fig. 1). Further assessment of genotype-specific differences was focused on the 6-month-old homozygous mice.

**Figure 2.**
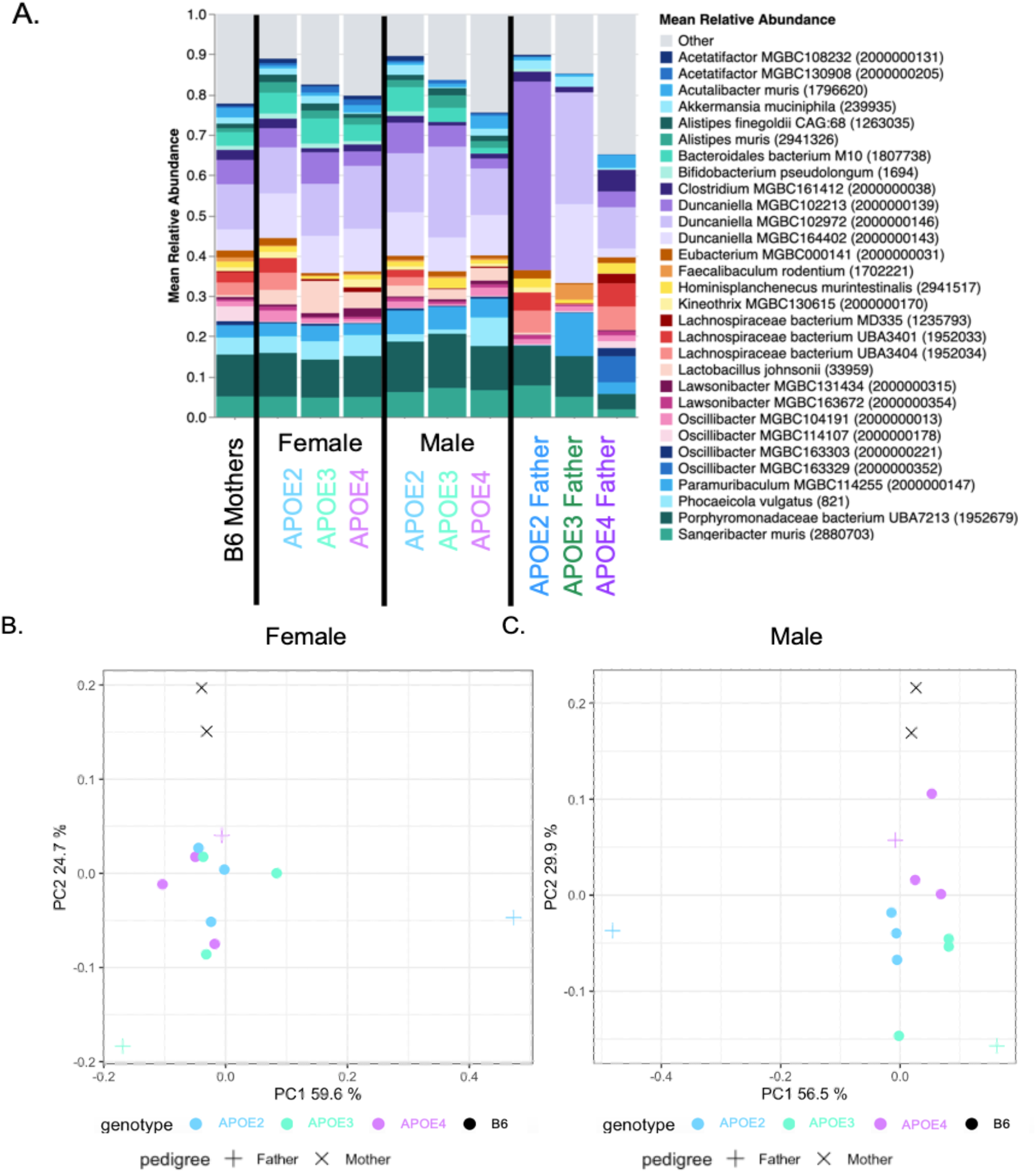
Microbiome standardization analysis of male and female heterozygous *APOE* 2,3, and 4 pups at wean. A. At wean top 30 gut microbiome bacterial species in stool. B. Gut microbiome principal component analysis (PCA) comparing relative abundance of species in stool among females and parents based on *APOE* genotype. C. Gut microbiome PCA comparing relative abundance of species in stool among males and parents based on *APOE* genotype. Stool was collected from the parents at breeder set up. n=3/sex/genotype for pups, n=1/genotype for fathers, n=2 for mothers.

Mean relative bacteria abundance and Principal Component Analysis (PCA) plots comparing homozygous pups at wean and 6-months-old with their grandfathers (purchased homozygous *APOE* 2, 3 and 4 males), demonstrated continued divergence of pups from their grandfathers (Sup. Fig. 2C-F). There were no significant differences in alpha diversity based on the Shannon index among *APOE* 2, 3, or 4 genotypes of heterozygous or homozygous female and male mice (Sup. Fig. 4). Several species of bacteria, especially species in the *Lachnospiraceae* family had significantly different abundances among homozygous *APOE* 2, 3, and 4 mice at 6 months of age (Fig. 3A and Sup. Fig. 5). GMB significantly differed by genotype at 6 months in homozygous females (p<0.001 by PERMANOVA), with *APOE*2 females demonstrating the greatest GMB divergence (Fig. 3B). GMB composition was also significantly different among genotypes in males (p<0.01 by PERMANOVA), but this was not as pronounced as the female mice divergence (Fig. 3C).

**Figure 3.**
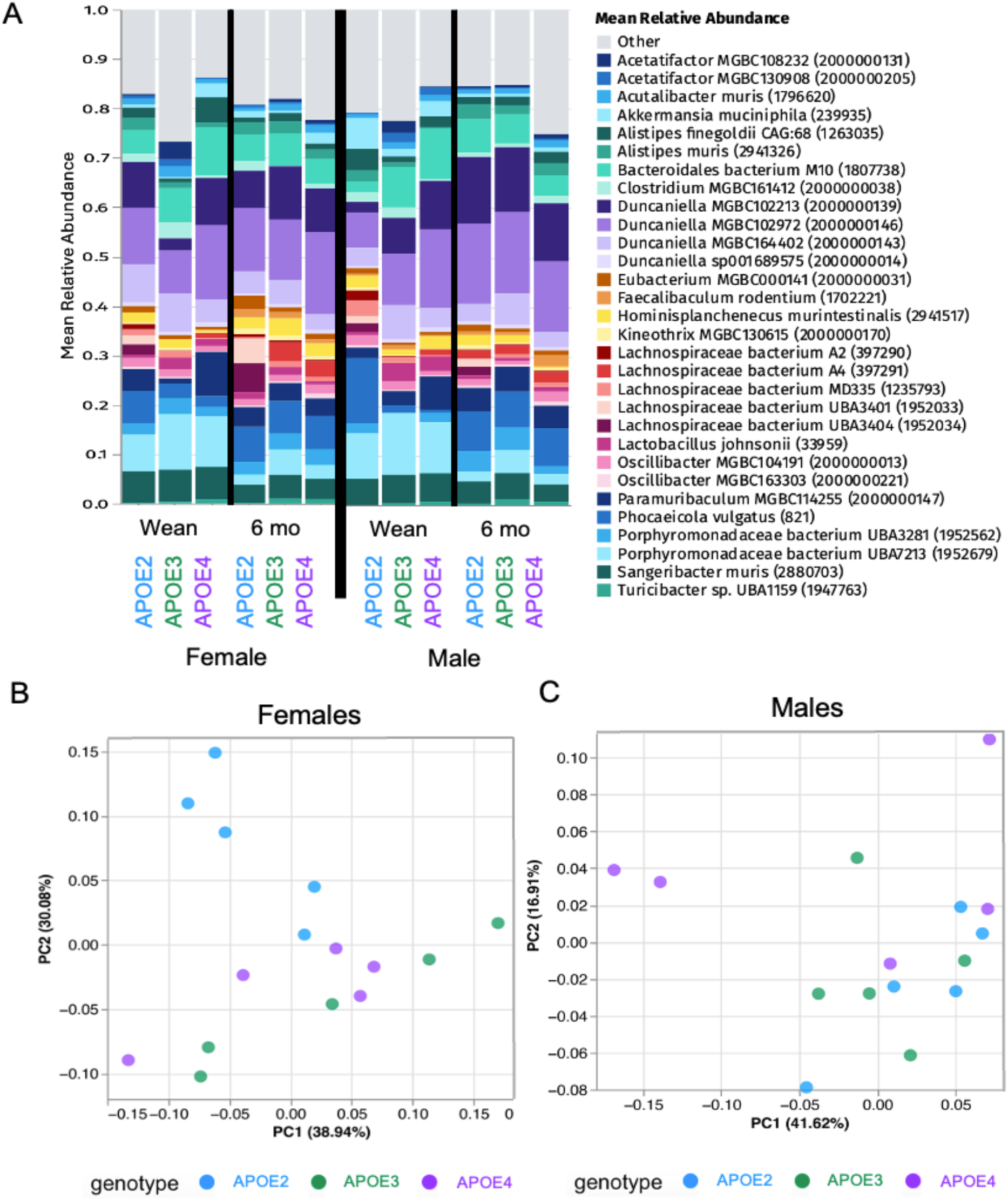
Genotype specific gut microbiome divergence of homozygous male and female *APOE* 2,3 and 4 mice. A. Top 30 gut microbiome bacterial species abundance in stool at wean and 6 months of age, respectively. C. and D. PCA plots comparing relative abundance of bacterial species in stool at 6 months of age of females and males, respectively. n=5/sex/genotype

To investigate microbiome differences between genotypes, linear discriminate analysis effect size (LEfSe) was used to indicate bacterial species that drive the differences in genotype composition by coupling statistical significance tests (p < 0.05 and fold change < 2) with biological consistency^26,27^. Cladograms demonstrate a visual representation of the LEfSe data showing phylogenic relationships of GMB by genotype for *APOE* 2, 3, or 4 homozygous and heterozygous 6-month-old female and male mice (Sup. Figs. 6 and 7, respectively). LEfSe and corresponding cladograms of 6-month-old homozygous *APOE* 2, 3, and 4 female mice demonstrate that *APOE4* females harbor the largest number of genotype-specific species, and *APOE3* females harbor species from the most distinct families (Fig. 4A, Sup. Fig. 6, and Sup. Table 1). *APOE*2 female-associated bacterial species were dominated by species within the *Lachnospiraceae* family (43% of species), and they had an overall higher abundance of *Lachnospiraceae* in comparison to their *APOE*3 and *APOE*4 female counterparts (p<0.01 and p<0.05, respectively) (Fig. 4A, Sup. Table 1, and Sup. Fig. 8A). Bacteria in the *Turicibacteraceae* family, a bacterial family associated with LOAD^28^, were only associated with *APOE3* mice, both males and females, and had the lowest overall abundance in *APOE*2 mice (Figs. 4A-B, Sup. Tables 1 and 2, and Sup. Fig. 8B). Among males, *APOE2* mice demonstrated the largest number of genotype-associated bacterial species, with *Lachnospiraceae* family representing 47% of the bacteria indicative of the *APOE2* genotype and increased *Lachnospiraceae* compared with their *APOE3* and *APOE4* counterparts (Fig. 4B, Sup. Figs. 6 and 8A, and Sup. Table 2). A coda4microbiome analysis, which is an updated approach to the Selective balance (Selbal) analysis, was performed to discover microbial signature predictive of genotype (Sup. Fig. 9)^29^. The coda4microbiome analysis uses a log-contrast model with positive values corresponding to group A (e.g. *APOE3*), negative values corresponding to group B (e.g. A*POE4*) and zero values corresponding to neither (not part of the microbial signature)^29^. Similar bacterial species were identified as indicative of each sex and genotype using the coda4microbiome analysis compared with the LefSe analysis.

**Figure 4.**
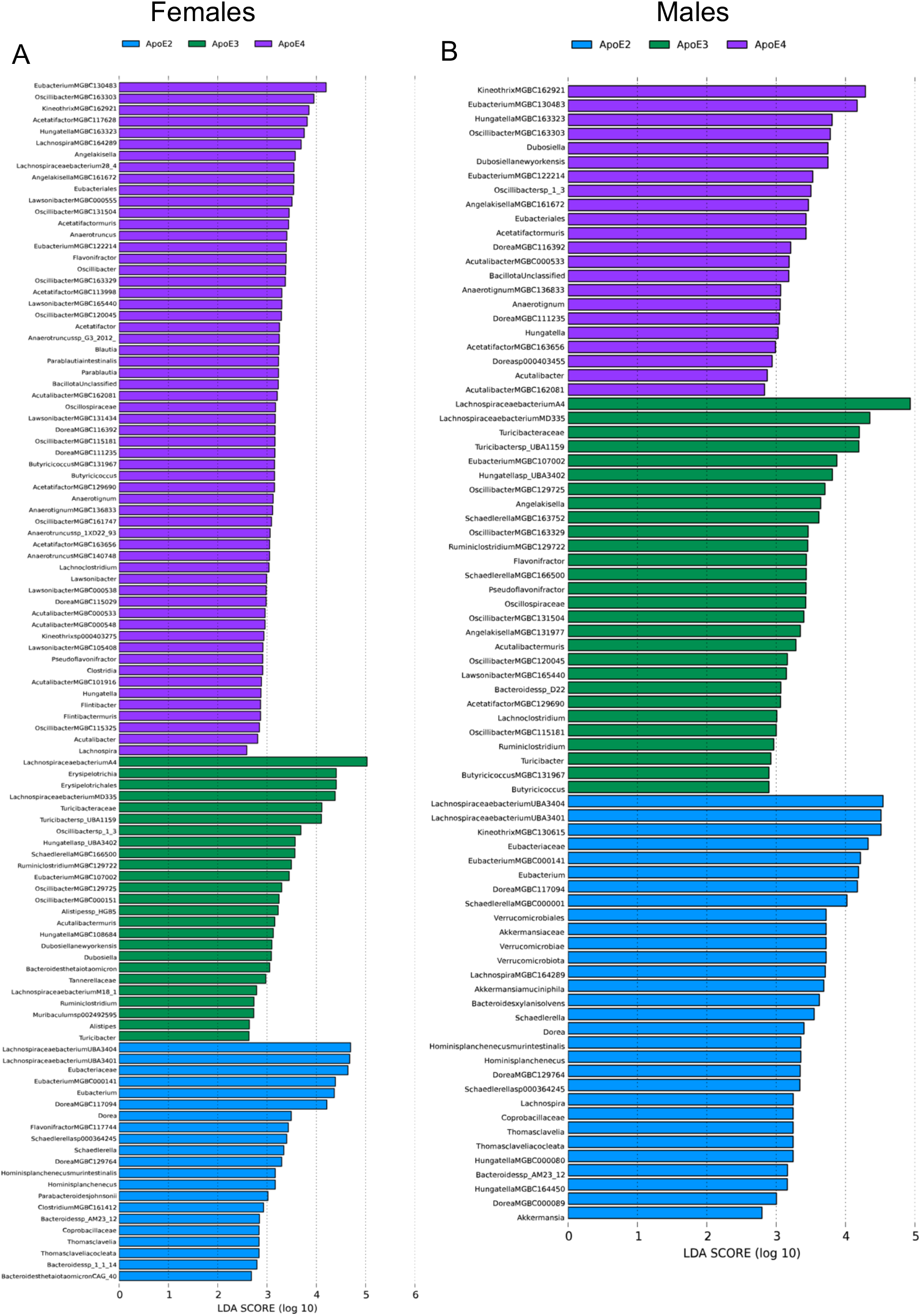
*APOE* genotype indicative bacterial species, genus, and families in 6-month-old homozygous *APOE* 2,3 and 4 mice. Linear discriminant analysis effect size (LEfSe). Only bacteria with an LDA score>2 and p<0.05 are shown. n=5/sex/genotype A. Females B. Males

## Discussion

The gut microbiome (GMB) aids in immune system development, is required for nutrition availability and uptake, involved in the host’s metabolism, and aids in the progression or delay of multiple diseases^2,3,14,30,31^. With the GMB being correlated with an ever-increasing number of diseases in the host, understanding how the GMB changes based on genotype and sex has the potential to aid in disease diagnosis, prevention, and treatment. GMB treatments for patients are already being given as fecal microbiota transplants (FMT) for Clostridium difficile (*C. diff*) infections and tested as an AD treatment^30,32^. There has been an increasing interest in formulating a microbiota pill instead of using FMT, as there have been cases where the FMT recipient died due to an unknown pathogenic species being present (immunocompromised patients are especially susceptible)^32^. Studies focusing on specific beneficial bacterial species and their relationship to genetic risk factors and disease are needed.

The GMB’s response to *APOE* genotype and sex has not been fully explored. Many factors affect the GMB, making human microbiome study correlations difficult, as seen by the disagreement within the literature relating bacterial families to disease progression, including AD and CVD studies^33–37^. Mouse model studies have the potential to eliminate most of the stochastic effects and confounding variables seen in human studies. Mouse studies may add further evidence to *APOE*-genotype specific GMB changes found in humans and associated diseases, as standardizing the microbiome and tracking genotype specific changes allows for the assessment of genotype as a driver of GMB differences correlative genotype studies^20,38^.

Standardization of the starting microbiome is important to truly assess genotype-specific GMB changes^20^. There are multiple methods that have been used to standardize the microbiome of mice, including co-housing, fostering pups together at birth (fostering), embryo transfer (ET), or breeding a male transgene mouse to a female wildtype mouse of the background strain (maternal inheritance)^20^. Microbiome standardization via co-housing experimental mouse lines together, despite grooming and coprophagia, has limitations. There is not a standardized method for when and how long to co-house mice and co-housing of non-littermates is challenging with male mice. Fostering pups at birth together with a foster female also has many challenges, as females can only care for a limited number of pups per cage, the timing must be exact so that all experimental lines are ready to transfer, the foster females must be available to receive foster pups, and there are challenges related to care and survival of fostered pups. ET involves implantation of embryos into a pseudo pregnant female and requires coordinated timing and costly surgeries. Maternal inheritance is considered by some as the standard for microbiome standardization, as it is easy, quick, and has been shown to be effective^20^. This study used maternal inheritance to standardize the GMB and generate experimental humanized transgene *APOE* 2, 3 and 4 mouse lines. The goal was to the eliminate stochastic changes inherent to independent mouse lineages and to study *APOE* genotype and sex-specific microbiome changes in a controlled setting with identical environments. Shallow shotgun sequencing was used to track GMB divergence (after GMB standardization) of humanized transgene *APOE* 2, 3, and 4 mice to 6 months of age down the species level.

In this study, humanized *APOE* 2, 3, or 4 male transgene mice (congenic on a C57BL/6J background) were purchased from JAX, and material inheritance was utilized. Humanized APOE 2, 3, and 4 transgene males were separately bred with the same two C57BL/6J sisters to standardize the microbiome of the pups. The resulting heterozygous *APOE* 2, 3, and 4 pup’s microbiomes were more similar to each other than their respective homozygous *APOE* 2, 3, or 4 fathers at wean. Sex and genotype divergence of the heterozygous pups by 6 months of age was observed without reversion to that of their fathers (2-3 months of age, JAX microbiome). A greater divergence was observed among females than males, with *APOE*2 females showing the greatest divergence. The homozygous progeny continued to display genotype and sex-specific bacterial GMB composition without reversion to their grandfathers (JAX microbiome), including differences in several bacterial species found in families that have been previously described in human AD patient studies as beneficial (*Lachnospiraceae*)^3,39,40^ or associated with poor outcomes (*Turicibacteraceae*)^28^.

Of the bacterial species demonstrated to be indicative of genotype in females, *APOE4* females had several butyrate/SCFA producers, including *Acetatifactor MGBC113998, Acetatifactor MGBC129690, Lawsonibacter MGBC000555, and Kineothrix MGBC162921*^41^, as well as *Lachnospiraceae bacterium 28-4*, which require SCFAs from other bacteria and is beneficial for cardiometabolic health^42^. *APOE4* females also had an increase abudance in *Kineothrix sp 000403275*, a bacteria recovered from inflamed mice, and *Acetatifactor muris*, a bacteria found in obese mice^43,44^. *APOE2* females had an increase in abundance of *Lachnospiraceae* species known to produce beneficial metabolites, including *Lachnospiraceae bacterium UBA3401*^20^. Finally, *APOE3* females had an increased abundance in species generally thought to be associated with health, including *Oscillibacter sp 1-3, Muribaculum sp 002492595*, and *Lachnospiraceae bacterium A4* and a bacterial species associated with anti-aging, *Dubosiella newyorkensis*^45–47^. However, different trends were observed in male mice. Although *APOE4* males demonstrated increases in *Acetatifactor muris*, similar to *APOE*4 females, they also demonstrated differences in the butyrate/SCFA producer *Kineothrix MGBC162921*, and health-associated bacteria, *Oscillibacter sp 1-3* and *Dubosiella newyorkensis*, similar to *APOE3* females^41,46,47^. *APOE3* males were associated with butyrate/SCFA producers, including *Acetatifactor MGBC129690* and *Lachnospiraceae bacterium A4*. Finally, *APOE2* males also demonstrated increases in specific butyrate/SCFA producers, namely *Kineothrix MGBC130615*, but also demonstrated an increase in *Muribaculum sp* 002492595 and Akkermansia muciniphila, bacteria associated with lower vascular inflammation^45,46^. *APOE*2 males associated with an increased abundance in bacteria with probiotic functions and a relationship to plant-rich diets in humans^32,41,43,47-49^. Many identified species have not been studied for their potential function and impact on health and disease. However, these results indicate some specific bacterial alterations may be investigated for their functional relevance to disease in humanized *APOE* transgene mice. In addition, several of these bacteria are also found in humans, including *Bacteroides xylanisolvens*, or have known human homologs, like *Dubosiella newyorkensis* in mice corresponding to *Clostridium inoculum* in humans^50^. Additional bacterial functional studies will provide acuity on metabolites affecting the host, the corresponding bacterial species in humans, and their interplay with other bacteria.

Shannon diversity did not show a significant difference in GMB species diversity between APOE genotypes. The greatest and fastest divergence was observed in *APOE*2 females with their high abundance of *Lachnospiraceae bacterium UBA3401* and *3404* as drivers of divergence (P<0.0001 for both). *Lachnospiraceae bacterium UBA3401* is considered a highly beneficial bacteria that has the potential help with clearance of *C. diff* infections via metabolites that decrease *C. diff* survival^32^. *APOE4* females were the only group associated with increases in bacteria from the genus *Blautia*, which are a known SCFA producers, and the only group with high levels of *Lachnospiraceae bacterium 28-4. Blautia*, although SCFA producers, are also associated with elevated blood pressure, and with a worse amyloid beta ratio in the CSF and can be an indicator of gut barrier integrity (GBI)^34,51,52^. Interestingly, *Lachnospiraceae bacterium 28-4* is known to increase in the presence of higher levels of SCFAs but not produce them^42^, further representing the interplay of metabolites of different bacterial species increasing or decreasing survivability of other bacteria. *APOE3* females and *APOE2* males demonstrated genotype-specific changes the widest variety of bacterial families, in comparison to their same sex counterparts (Fig. 4). Previous human, mouse, and bacterial studies consider a few bacterial families to be generally beneficial for everyone, including members of the *Lachnospiraceae* family and SCFA producers, which were also found to be altered in our study^31,32,35,39,40^.

It is known that sex hormones affect the GMB, genotype, and disease progression. Here, we demonstrate that *APOE* genotype affected females more than males, and that some bacterial species indicative of one genotype in females, was indicative of a difference genotype in males (Fig. 5, Supp. Tables 1 and 3). Many studies that identify bacteria at the family or genus level disagree with which bacteria are indicators of known diseases, which is likely the result of sex, GMB present at the start of the experiment, and the importance of specific species that may shift the survivability of other bacteria in the presence or absence of a particular gene^33–37^. Studies that identify bacteria at the species level will provide better insights into the microbial composition and the interplay between bacterial species, genetics, and disease.

**Figure 5.**
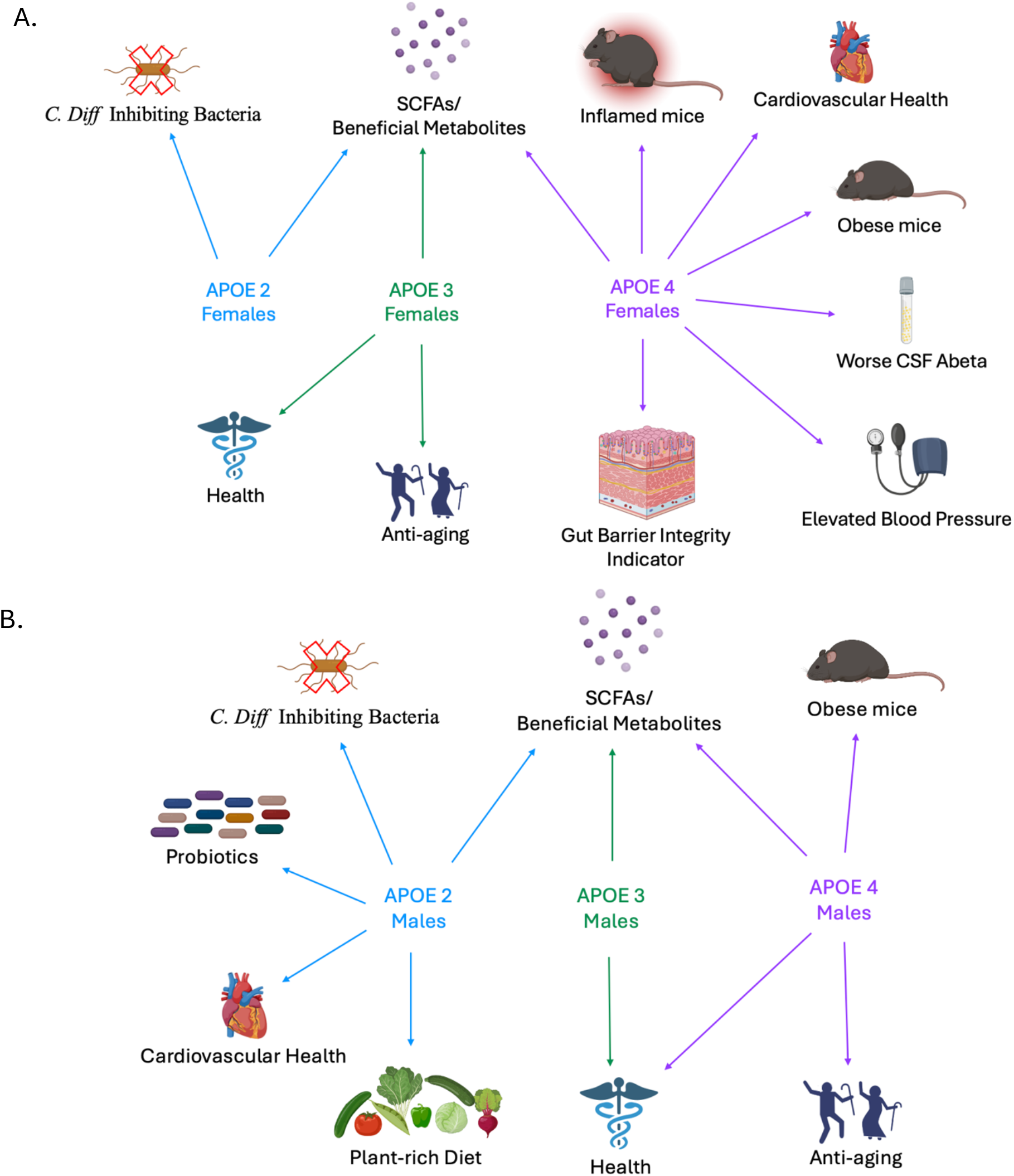
Interplay of APOE genotype, sex, and gut microbiota. Known associations of bacteria and their metabolites found in homozygous 6-month-old humanized APOE 2, 3, and 4 mice. A. Females B. Males

Overall, *APOE2* females and males demonstrated an increased quantity of anti-inflammatory profile associated bacteria including in the family *Lachnospiraceae (Lachnospiraceae bacterium UBA3401, 3404*)^20^ and decreased *Turicibacteraceae* which has been associated with AD in previous human studies^28^ in comparison to their *APOE3* and *APOE4* counterparts. Some known SCFA producing bacterial species varied by *APOE* genotype, with different species associating with different genotypes. Although the number and overall quantity of bacteria within the family *Clostridiaceae* did not change by genotype, bacteria in the *Blautia* genus were only associated with APOE4 females, demonstrating a need for species level resolution.

This study has several limitations. The heterozygous and early homozygous timepoints include only 3 samples per sex per genotype, which limits the conclusions that can be drawn over time. In addition, samples were sequenced in batches, which required rarefication to standardize the data (sum scale method). However, batch effects after rarefication were assessed and none were found. The pups that were homozygous in the N1F1 generation could not be used as the homozygotes in this study, because they were used as breeders to generate our homozygous microbiome standardized colony. Moreover, breeding likely changed the N1F1 generation microbiome. Therefore, this study investigated the N1F2 generation, which is one generation removed from standardization. However, data from 6-month-old heterozygotes demonstrated similar trends and supports the genotype specific differences we observed (Sup. Figs. 1-3). Finally, one key limitation as is the current lack of knowledge available on specific bacterial species, and future studies should investigate the functional differences of GMB composition following deeper sequencing.

Studies exploring inflammatory profiles of prevalent bacterial species to better understand how they interact with the host and each other, and how they are affected by host genes are needed. The *APOE*-specific bacterial species’ impact on inflammatory responses *ex vivo* and i*n vivo* should be studied to further untangle the complexities of *APOE*-dependent GMB composition, as well as longevity and robustness of FMT or specific species and strain level microbiome supplementation as an option to adjust the GMB of *APOE4* individuals. This study shows the first data demonstrating GMB divergence over time driven by *APOE* genotype from an inherited source. Studying genotype specific changes in GMB and how specific *APOE*-specific bacteria impact inflammatory responses can aid in understanding and potentially treating some of the underlining associated diseases.

## Abbreviations

AD: Alzheimer’s disease
ApoE: Apolipoprotein E protein
*APOE*: Apolipoprotein E gene
APOE 2, 3, or 4: Humanized mice with either *APOE* 2, 3, or 4
APOE 2: Humanized APOE 2 mice
APOE 3: Humanized APOE 3 mice
APOE 4: Humanized APOE 4 mice
B6: C57BL/6J
CVD: Cardiovascular disease
GMB: The gut microbiome
JAX: The Jackson Laboratory
LefSe: Linear discriminate analysis Effect Size
LOAD: Sporadic or late onset Alzheimer’s Disease
PCA: Principal component analysis
SCFAs: Short chain fatty acids

## Ethical Statement

All animal studies were conducted at the McLaughlin Research Institute (MRI) between September 2022 and March 2024. The MRI Institutional Animal Care and Use Committee (IACUC) approved all protocols. All procedures and protocols in this study were performed in accordance with the MRI IACUC protocols, the NIH *Guide for the Care and Use of Laboratory Animals* (National Academies Press, 2011), the ARRIVE guidelines (PMID 32663221), and the *Public Health Service Policy on Humane Care and Use of Laboratory Animals* (Public Health Service, 2015). MRI is an American Association of Laboratory Animal Science (AALAS) certified institute.

## Author Contributions

MAM assisted with experimental procedures and design, conducted the study, conducted data analysis, conceptualized, drafted, and edited the article. HG conducted the study, conducted data analysis, drafted, and edited the article. MM conducted the study. FRC conducted data analysis and wrote associated methods. AG conducted the study and assisted with the article. AP conducted data analysis. THM conceptualized and designed the experiment, conceptualized, drafted, and edited the article.

## Data Availability

The raw microbiome data files will be deposited in an NIH approved publicly available database once the paper has been accepted for publication, and this section will be edited to include the database name and the Digital object identifiers (DOIs) or accession numbers (add URLs) for the final draft.

## Acknowledgements

The authors would like to thank colleagues and staff at the McLaughlin Research Institute for their support of these studies. Part of Figures 2 and 5 were made using BioRender.

